# Human regulatory T-cells locally differentiate and are functionally heterogeneous within the inflamed arthritic joint

**DOI:** 10.1101/2022.02.18.480998

**Authors:** Lisanne Lutter, M. Marlot van der Wal, Eelco C. Brand, Patrick Maschmeyer, Sebastiaan Vastert, Mir-Farzin Mashreghi, Jorg van Loosdregt, Femke van Wijk

## Abstract

**Objective:** Tregs are crucial for immune regulation, and environment-driven adaptation of effector (e)Tregs is essential for local functioning. However, the extent of human Treg heterogeneity in inflammatory settings is unclear.

**Methods:** We combined single-cell RNA- and TCR-sequencing on Tregs derived from 4-6 patients with juvenile idiopathic arthritis (JIA) to investigate the functional heterogeneity of human synovial fluid (SF)-derived Tregs from inflamed joints. Confirmation and suppressive function of the identified Treg clusters was assessed by flow cytometry.

**Results:** Four Treg clusters were identified; incoming, activated eTregs with either a dominant suppressive or cytotoxic profile, and GPR56^+^CD161^+^CXCL13^+^ Tregs. Pseudotime analysis showed differentiation towards either classical eTreg profiles or GPR56^+^CD161^+^CXCL13^+^ Tregs supported by TCR data. Despite its most differentiated phenotype GPR56^+^CD161^+^CXCL13^+^ Tregs were shown to be suppressive. Furthermore, BATF was identified as an overarching eTreg regulator, with the novel Treg-associated regulon BHLHE40 driving differentiation towards GPR56^+^CD161^+^CXCL13^+^ Tregs, and JAZF1 towards classical eTregs.

**Conclusion:** Our study reveals a heterogeneous population of Tregs at the site of inflammation in JIA. SF Treg differentiate to a classical eTreg profile with a more dominant suppressive or cytotoxic profile that share a similar TCR repertoire, or towards GPR56^+^CD161^+^CXCL13^+^ Tregs with a more distinct TCR repertoire. Genes characterizing GPR56^+^CD161^+^CXCL13^+^ Tregs were also mirrored in other T-cell subsets in both the tumor and autoimmune setting. Finally, the identified key regulators driving SF Treg adaptation may be interesting targets for autoimmunity or tumor interventions.

## Introduction

Regulatory T-cells (Tregs) comprise a subset of CD4^+^ T-cells crucial in preserving immune homeostasis by antagonizing immune responses. The transcription factor (TF) FOXP3 characterizes Tregs, and mutations in the *FOXP3* gene lead to severe inflammation in both mice and humans^1,2^. In recent years, potential therapeutic strategies targeting Tregs in both the autoimmune and tumor setting have been explored. In autoimmunity the number and/or functionality of Tregs should be enforced, whereas in the tumor milieu the suppressive capacity of Tregs should be dampened^3,4^. This can, amongst others, be achieved by expanding Tregs, or by blocking Treg functioning via immune checkpoint blockade. PD-1 and CTLA-4 blockade are employed in the cancer setting and several other targets are currently tested in clinical trials^4^. Understanding the heterogeneity and plasticity of Tregs is essential in understanding immunodynamics at play in health and disease. This knowledge can then be exploited to develop and improve (potential) Treg-based therapeutic strategies.

Tregs are not identical in every tissue of residence, but are tailored to the environment in which they have to function, and can alter their phenotype according to micro-environmental changes over time^5,6^. This plasticity enables continuous adaptation to changing immunodynamics. Upon activation, Tregs gain an effector profile (eTreg) and can initiate transcriptional programs associated with the dominant T helper (Th) response at the site of inflammation to enable Treg survival and suppression of the respective Th-cells^5,7,8^. These co-transcriptional programs include Th1 (T-bet), Th2 (GATA-3), Th17 (RORc), and T follicular regulatory cell (Tfr, Bcl6) programs^9^. We have recently demonstrated that in synovial fluid (SF) of patients with juvenile idiopathic arthritis (JIA), a predominantly Th1/Th17-associated disease^10^, Tregs retain a functional Treg core profile and obtain a Th1-skewed co-transcriptional profile on the proteomic, transcriptomic and epigenetic level^7^. Furthermore, it has been shown that some Tregs can acquire additional functions including stimulation of tissue repair in the intestine^11^ and hair follicle growth in the skin^12^. These additional functions further elucidate the local function and relevance of Tregs. An outstanding knowledge gap regarding human Tregs is the degree of heterogeneity of Tregs present within inflammation, if heterogeneity has consequences for Treg function, and how this relates to clonality and thus the T-cell receptor (TCR)-repertoire of Tregs.

Tregs in tissues and peripheral blood (PB) have been shown to be heterogeneous^5,7,8^, but data on local inflammatory environments in humans are lacking. Single cell (sc)RNA-sequencing enables us to map cellular states within an environment to facilitate our understanding of the phenotypic plasticity and functional diversity within the Treg population. SF-derived Tregs from JIA patients provide us with a relevant model of local autoimmune inflammation to determine the functional differentiation and clonality of inflammation-derived Tregs and its regulators. In this study we aimed to dissect the heterogeneity of SF-derived Tregs by employing scRNA-sequencing to further elucidate the immunodynamics at play in an inflammatory environment, specifically in JIA.

## Results

### Heterogeneity within inflammatory synovial fluid Tregs

To assess the heterogeneity at the site of inflammation in JIA patients, SF Tregs (live CD3^+^CD4^+^CD127^low^CD25^high^) from three patients with oligoarticular JIA were sorted for single cell transcriptome analysis. Dimensionality reduction of 980 Tregs after quality control revealed presence of four clusters within the Treg population (Figure 1a), with each cluster present in all three patients (Figure 1b). All clusters were of Treg origin with the vast majority of cells (97%) expressing at least one FOXP3 mRNA molecule and in 99.96% of the Tregs the human core Treg signature as defined by Ferraro *et al*.^13^ was enriched (Supplementary figure 2b). There was no cluster-specific association with the cell cycle phase the Tregs resided in (Supplementary figure 2c; Pearson’s Chi-squared test, *p* = 0.8512).

**Figure 1.**
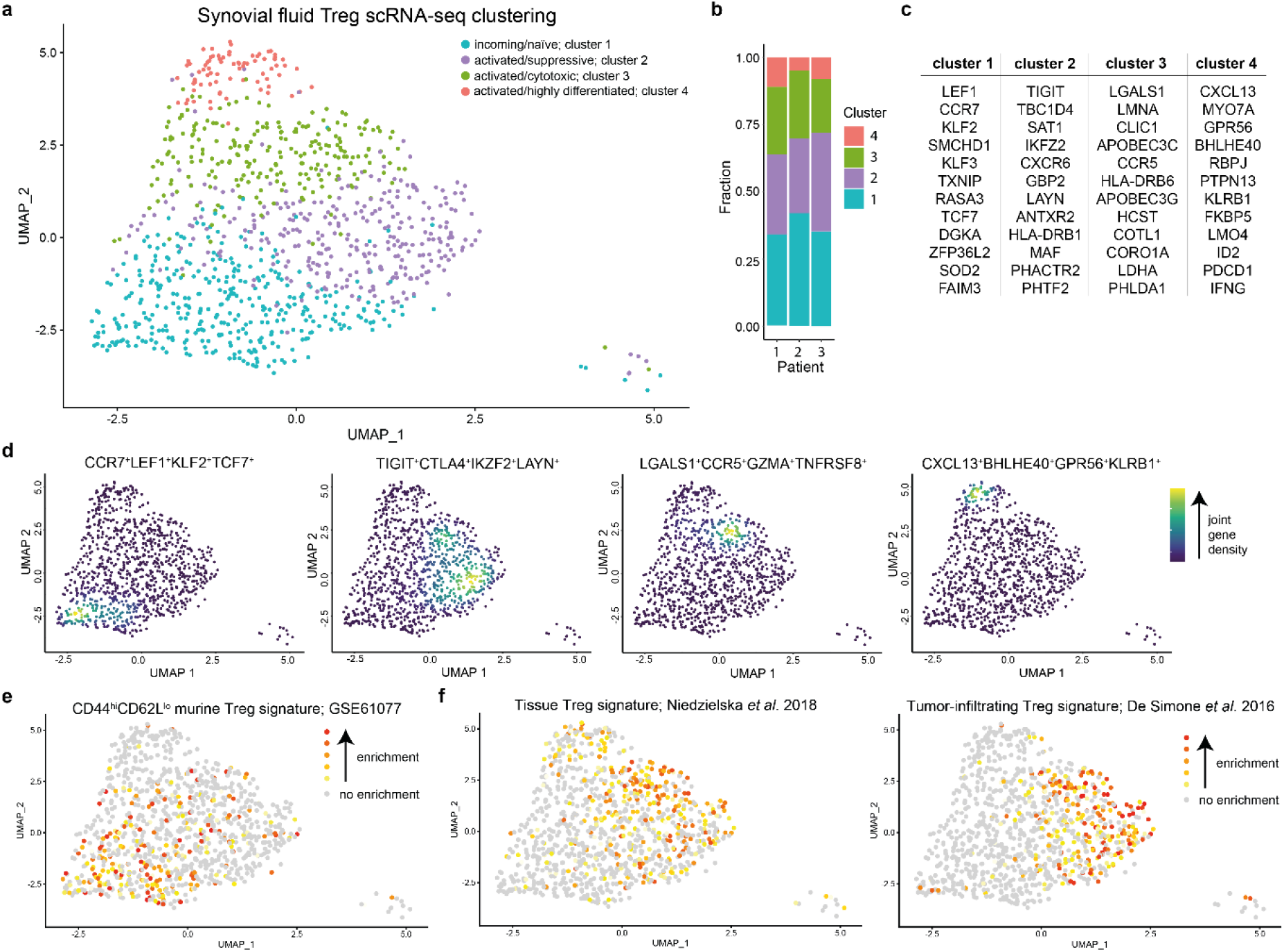
Heterogeneity and phenotypical profile of synovial fluid Tregs. **(a)** Dimensionality reduction (UMAP) of all synovial fluid (SF)-derived Tregs (sorted on live CD3^+^CD4^+^CD127^low^CD25^high^) of three Juvenile Idiopathic Arthritis (JIA) patients. Tregs are colored based on the assigned cluster. **(b)** Reproducible composition of the Tregs across the three included patients. Y-axis: fraction of cells colored based on the cluster as shown in **(a)** and separated per patient on the x-axis. **(c)** Top 12 upregulated genes, based on *p*-adjusted value, per cluster based on MAST differential gene expression analysis. **(d)** UMAPs of the combined expression of 2-4 selected differentially expressed genes per cluster shown in nebulosa density (kernel density estimation to handle sparsity of scRNA-sequencing data). The scale ranges from blue to yellow, with in yellow the highest kernel density, thus the highest (estimated) expression of all combined selected genes. **(e)** Gene set analysis of a gene module downregulated in naive versus memory CD4 T-cells (GSE61077). Enrichment of a gene set is calculated per cell; grey signifies no enrichment of the gene set and yellow to red represents increasing enrichment. **(f)** Similar to **(e)**, but for a human shared tissue Treg signature ^56^ and a human tumor-infiltrating Treg signature^57^.

The largest cluster (37.24%), cluster 1, was characterized by genes that are downregulated upon activation and maturation of T-cells, including *CCR7, LEF1, KLF2, KLF3* and *TCF7*, suggesting a relatively quiescent/resting phenotype and probably representing Tregs that only recently migrated into the inflamed joint. The other three Treg clusters all displayed an activated gene signature including expression of many MHC class II genes (e.g. *HLA-DR, -DQ, -DP, -DM)*, but also *DUSP4, CTSC, CTSB, ITM2A* and *LMNA* amongst others). Additional markers separated these clusters from each other. Both cluster 2 (31.22%; *TIGIT, CTLA4, IKZF2, LAYN*) and 3 (23.57%; *LGALS1, CXCR6, CCR5, TNFRSF8, GZMA*) showed high expression of genes associated with highly suppressive Tregs. The smallest activated Treg cluster, cluster 4 (7.96%), expressed a set of genes not commonly or previously associated with Tregs (*CXCL13, GPR56, MYO7A, BHLHE40, PTPN13, KLRB1*) (Figure 1c, d, Supplementary figure 2d, 3, Supplementary table 1). Overall, the three activated Treg clusters showed expression of a wide array of co-stimulatory and co-inhibitory markers to help suppress immune activation in a mostly contact-dependent manner but with a per cluster different potential dominant mode of suppression (e.g. *CTLA4* in cluster 2, *GZMA* in cluster 3, *LAG3* in cluster 4; Supplementary table 1).

Area under the curve (AUC) analysis to determine whether a gene set is active in a cell^14^ supported the observation that cluster 1 comprises primarily resting Tregs with genes being upregulated in naive compared to memory Tregs (39.2% enriched Tregs in cluster 1 compared to 19.3%, 14.3% and 5.2% for cluster 2-4; Pearson’s Chi-squared test *p* = 2.2×10^−16^) or naive compared to effector memory CD4 T-cells (37.6% of enriched Tregs in cluster 1 compared to 28.2%, 7% and 11.6% for cluster 2-4; Pearson’s Chi-squared test *p* = 2.2×10^−16^) (Figure 1e, Supplementary figure 2e). Gene signatures associated with eTregs as found in homeostatic tissues (37.3-41% enriched Tregs in cluster 2-4 compared to 18.1% for cluster 1; Pearson’s Chi-squared test *p* = 2.502×10^−10^) and the tumor tissue micro-environment (43.8/29.5% enriched Tregs in cluster 2/3 compared to 6.4/11.5% for cluster 1/4; Pearson’s Chi-squared test *p* = 2.2×10^−16^) were highly enriched in cluster 2 and 3 Tregs (Figure 1f). Subsequent gene ontology analysis also revealed that clusters 2-4 share upregulation of Th differentiation-associated genes, whereas cluster 1 showed downregulation of TCR-signaling pathways compared to clusters 2-4 indicative of resting Tregs. Additionally, cluster 3 and 4 shared pronounced upregulation of TCR- and Notch signaling, and of all SF-derived Tregs those from cluster 3 seemed to rely most on glycolysis (Supplementary table 2).

Microenvironmental cues can shape the transcriptomic signature of Tregs with a resulting co-transcriptional Th-cell program^5,7^. These can be distinguished based on the expression pattern of the chemokine receptors CXCR3, CCR4, CCR6, CCR10 and CXCR5^15^. CXCR5, linked to Tfr, was not expressed in any of the SF Tregs. Cluster 1 harbored Tregs expressing mixed chemokine receptor profiles associated with Th2, Th22, Th1 and Th17 cells although overall expression of CCR4, CCR6 and CCR10 was low. In contrast, cluster 2-4 Tregs were predominantly CXCR3^+^ (51.7%, Th1-associated) (Supplementary figure 4). These data indicate that Tregs in the inflammatory SF environment are heterogeneous, and that local cues preferentially induce a Th1 co-transcriptional program in these Tregs.

### Clonotype sharing amongst synovial fluid-derived Treg clusters

Next, we assessed whether the TCR of individual Tregs skews differentiation to a certain phenotype upon triggering. Therefore, we employed a 10X Genomics dataset, including both 5’ gene expression and TCR sequences, published by Maschmeyer *et al*.^16^ containing SF-derived Tregs from JIA patients. Dimensionality reduction revealed similar Treg clusters as in our dataset indicating that the clustering is robust (Supplementary figure 5a).

On average, 55% of the detected clonotypes (full length combined TRA and TRB chain) were unique within a patient (range 34.8-72.7%), showing presence of clonal expansion. Exploratory analysis revealed that cluster 1 contained mostly single frequency clonotypes (average of 65.7% single clonotypes, range of 42.0-89.6%), whereas for cluster 2-4 this was less (average of 49.2%, 44.3% and 61.4%, respectively) (Figure 2a). Clonal expansion was analyzed by dividing the detected clonotypes in 6 groups based on frequency. The proportional space filled by the most expanded clones revealed a gradient from cluster 1 to 4. In cluster 4, the most prevalent clonotypes comprised the greatest proportion of the total clones present (Figure 2b). A wide spread was observed for cluster 1 for the number of clonotypes comprising 10% of the total repertoire with an average of 16 clonotypes (Figure 2c). For cluster 2 and 3 only five clonotypes and for cluster 4 merely two different clonotypes were observed on average (Figure 2c). Several diversity indices indeed indicated that cluster 1 had the most diverse clonal repertoire compared to the three activated Treg clusters (Supplementary figure 5b).

**Figure 2.**
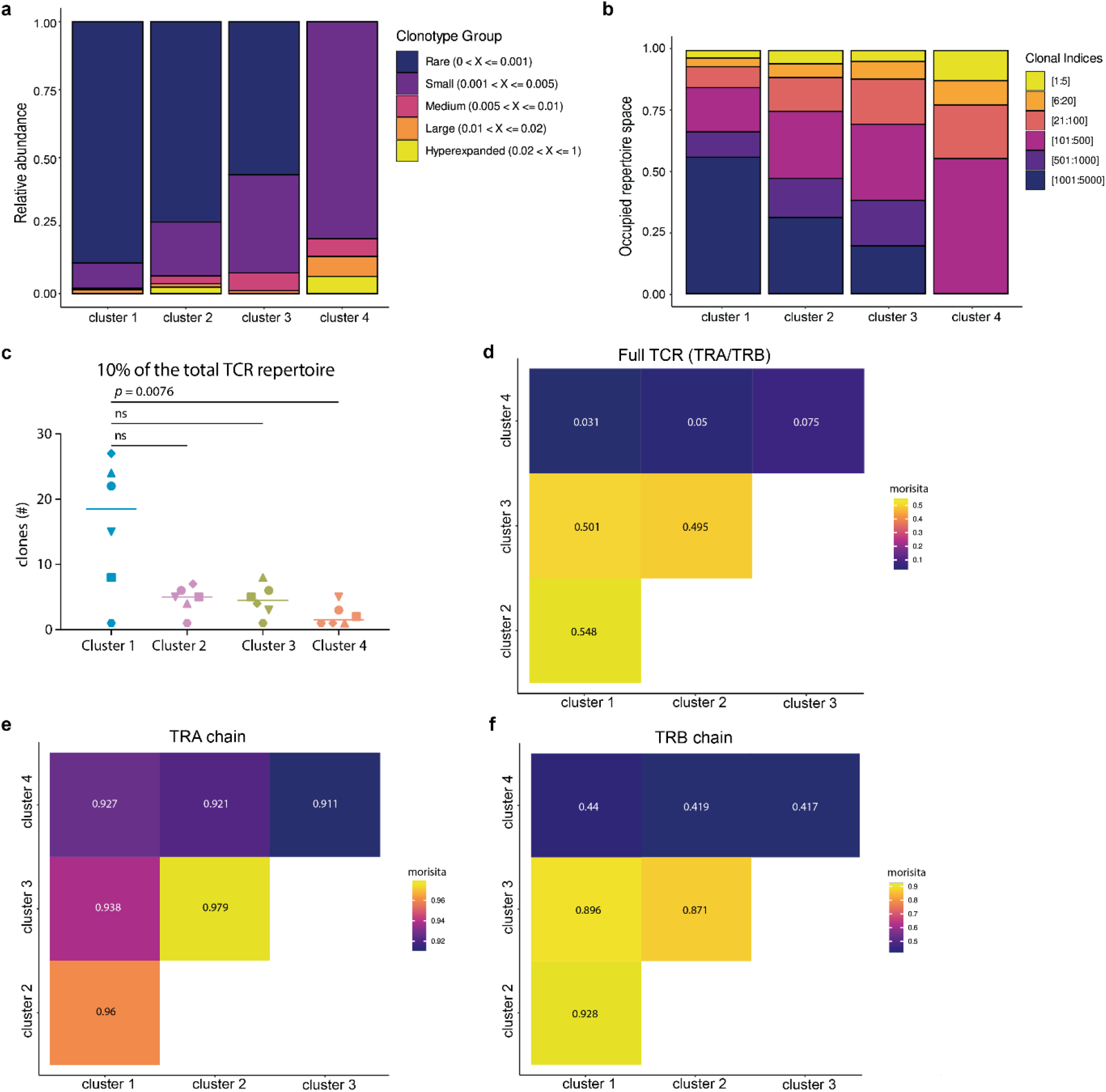
Clonal expansion and clonal overlap of synovial fluid-derived Tregs. **(a)** Histograms of the relative proportion filled up by clonotypes that form 1% (rare), 1-5% (small), 5-10% (medium), 10-20% (large) or > 20% (hyperexpanded) of the total repertoire, separated by cluster. 1% refers to clones that are present once (not expanded). **(b)** Histograms with the clones divided in 6 bins based on the frequency each clonotype is present. The bins are the top 1:5 clonotypes, 6:20, 21:100, 101:500, 501:1000, and 1001:5000. **(c)** Graph showing the least number of clonotypes (combined TRA and TRB chain) that comprise 10% of the TCR repertoire per patient (*n* = 6, different symbols) per cluster (colors as per Supplementary figure 5a). Comparison was performed with a Friedman’s test followed by Dunn’s post hoc analysis. **(d)** Morisita diversity index for the similarity of the TCR repertoire between all four SF Treg clusters based on the nucleotide sequence of both the TRA and TRB chain. The scale ranges from 0 to 1, with 0 indicating no overlap and 1 indicating identical repertoires. **(e)** Similar to **(d)** but for the nucleotide sequence of the TRA chain. **(f)** Similar to **(d)** but for the nucleotide sequence of the TRB chain.

We also determined presence of clonal sharing between the clusters, indicative of a shared origin. The overlap coefficient was calculated based on overlap of the complete TRA/TRB nucleotide sequences. This showed a clonotype overlap of 25% and 32.4% of cluster 1 with clusters 2 and 3, respectively. The latter two clusters shared 32.3% of their clonotypes. However, interestingly there was little clonotype sharing on nucleotide level (< 15%) between cluster 4 and the other clusters, although on TRA/TRB chain level this was 21.1-28.5%. Overall, based on the Morisita similarity index, a measure that takes the total number of cells into account, it was indeed clear there was little similarity between clusters 1-3 and cluster 4 Tregs (Figure 2d). For the TRA chain alone the similarity between all four clusters was very high suggesting that the differences are primarily formed by the TRB (Figure 2e). These data indicate that cluster 2 and 3 eTregs are relatively similar, as also suggested by their transcriptomic profile, whereas cluster 4 Tregs contain more distinct clonotypes primarily skewed by the TRB.

### Non-linear differentiation of Tregs within synovial fluid

To further explore how the SF Treg clusters are related, we employed pseudotime analyses to estimate Treg differentiation within SF based on transcriptional similarities. We applied Monocle v3^17,18^ to perform trajectory inference (Figure 3a) and pseudotime plotting (Figure 3b). Mathematical assessment of the potential starting node for the differentiation trajectory pointed towards cluster 1 Tregs. Those Tregs were indeed highest in (combined) expression of the genes *CCR7, LEF1, TCF7* and *KLF2* associated with a naive state (Figure 3c). The identified trajectory suggests that upon arrival within the SF environment, Tregs are skewed towards a classical eTreg phenotype (clusters 2 and 3) or towards cluster 4 Tregs, although cluster 4 Tregs may also pass a cluster 3 phenotype along the differentiation trajectory (Figure 3a, b).

**Figure 3.**
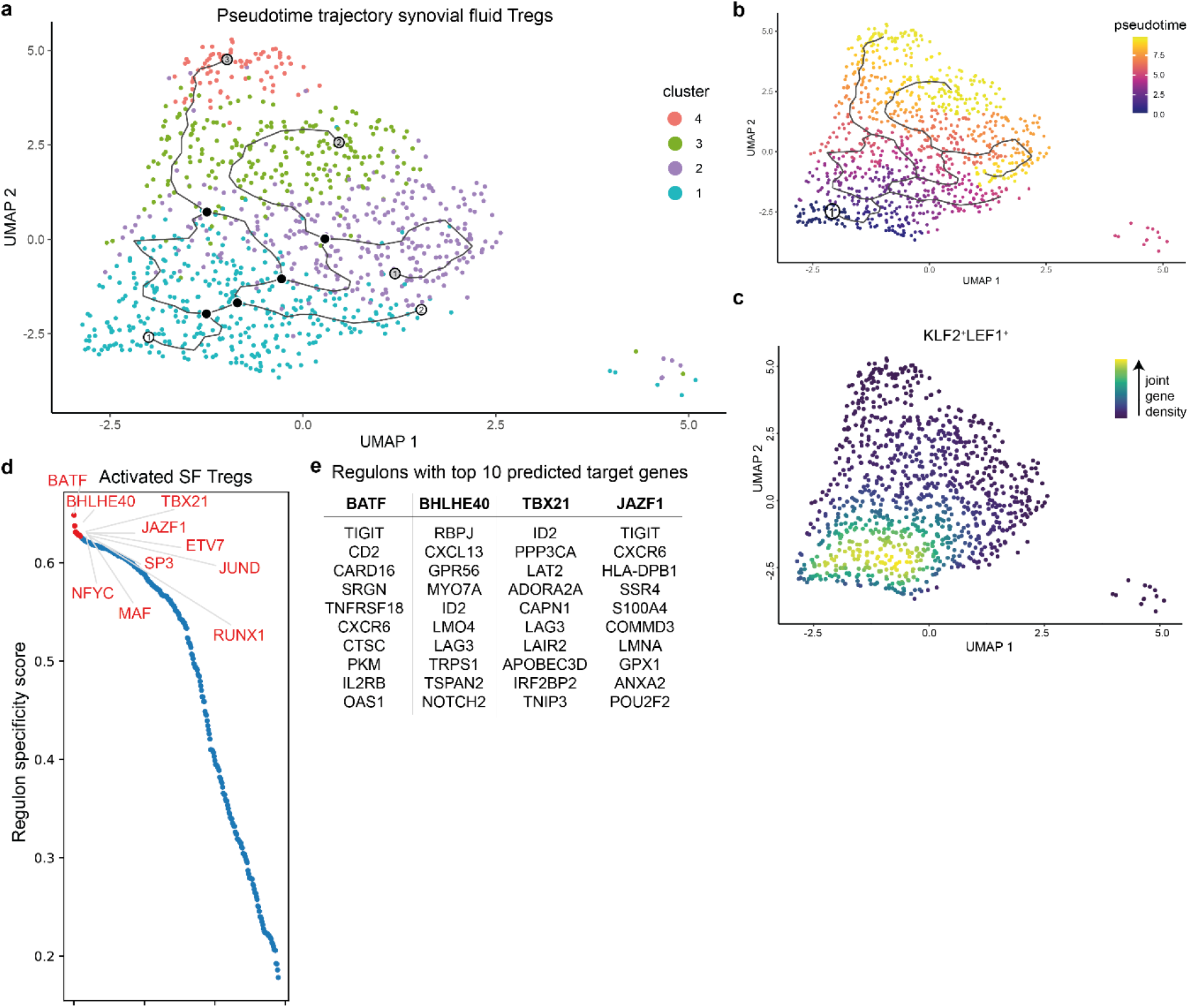
Predicted differentiation trajectory and regulation of synovial fluid Tregs. **(a)** Predicted differentiation trajectory of Tregs within the synovial fluid (SF) environment plotted on the UMAP as per Figure 1a. White circles represent predicted starting points, black circles designate decision points setting a cell upon a trajectory, and grey circles (ending in all three activated Treg clusters) depict end points. **(b)** Similar to **(a)** but colored based on pseudotime with early cells in blue and end stage cells in yellow. The circle with number 1 is the mathematically determined starting node. **c** UMAP of the combined expression of *CCR7, LEF1, KLF2* and *TCF7* in nebulosa density. The scale ranges from blue to yellow, with the highest kernel density displayed in yellow, thus representing the highest (estimated) expression of all combined selected genes. **(d)** Top 10 predicted regulons (transcription factor and its target genes) to drive differentiation of cluster 1 to activated cluster 2-4 (e)Tregs. The regulons are ranked by the regulon specificity score for cluster 2-4 Tregs shown on the y-axis ranging from 0 to 1, with 1 indicating complete specificity of the regulon for the cell type. **(e)** Selected regulons with their top 10 calculated target genes.

TFs interact with cis-regulatory elements and regulate cell differentiation, and together with their target genes it forms a regulon. SCENIC can be employed for gene regulatory network analysis to deduce active regulons per cell. Here, we compared cluster 1 recently migrated Tregs with cluster 2-4 activated (e)Tregs. There was no clear binarization of the regulons per cluster indicating that there is a gradual differentiation of SF Tregs. The top regulons were all upregulated in cluster 2-4 Tregs and associated with differentiation or maintenance of the Treg phenotype indicating that upon migration to the SF-environment cluster 1 Tregs differentiate towards cluster 2-4 Tregs and not in the opposite way (Figure 3d). BATF is a known key driver in eTreg differentiation^7,19,20^, and indeed, our analysis revealed BATF as the primary local regulon for differentiation of Tregs that recently migrated to SF. RUNX1 and NFYC have also been previously associated with Treg development, maintenance and differentiation, and TBX21 is a possible driver of the Th1-associated co-transcriptional program^7,21,22^, whereas for example SP3 is associated with generic processes including proliferation, apoptosis and metabolism^23^. BHLHE40 was identified as a novel regulon for eTreg differentiation, and target genes mostly comprised cluster 4-associated genes including *CXCL13, GPR56* and *KLRB1*. However, also amongst its target genes are genes required for Treg differentiation and survival such as *ID2*^24^. Other BHLHE40 associated genes, such as, *RBPJ* and *RFX1* have been linked to inhibiting Th2 and Th17 differentiation respectively^25,26^. Another novel identified regulator, i.e. JAZF1, primarily drives SF Treg differentiation towards cluster 2-3 Tregs. Its target genes include the (e)Treg genes *TIGIT* and *CXCR6*^6,7^, but also *POU2F2* which regulates a shift towards aerobic glycolysis^27^ (Figure 3e). Overall, these data indicate that Tregs arriving in the inflamed SF-environment can differentiate into activated (e)Tregs via different routes in a non-linear fashion, facilitated by key regulators including BATF, TBX21, RUNX1, and the newly (e)Treg-associated regulons BHLHE40 and JAZF1.

### GPR56^+^/CD161^+^/CXCL13^+^ synovial fluid Tregs are highly differentiated and suppressive

Cluster 4 Tregs were defined by genes not commonly associated with Tregs which warranted further investigation. On protein level, we could confirm presence of Tregs expressing GPR56 and/or CD161 (average of 18.1%, range 10.3-32.3%). This subset was specifically enriched for CXCL13 expression. In PB Tregs, expression of GPR56 and/or CD161 averaged 4.6%, with no CXLC13^+^ Tregs present (Figure 4a). In line with the transcriptomic data, GPR56^+^CD161^+^CXCL13^+^ SF Tregs expressed high levels of PD-1 and LAG3 (Figure 4b). We also confirmed FOXP3 protein expression within these Tregs. Although the intensity (median fluorescence intensity) of FOXP3 expression in GPR56 and/or CD161 positive CXCL13^+^ SF Tregs was lower compared to other SF Tregs it was higher than in PB Tregs and PB and SF non-Tregs (Figure 4c). In addition, a gradual decline of Helios-expressing Tregs was observed from PB Tregs to CXCL13^+^ SF Tregs; however, compared to non-Tregs about 5.5x more cells express Helios (53.9% for CXCL13^+^ SF Treg versus 9.9% for CXCL13^+^ SF non-Treg) (Figure 4d). Even though Helios cannot distinguish thymic- and peripherally-derived Tregs, its expression in FOXP3^+^ Tregs indicates a stable and activated phenotype^28,29^, suggesting that CXCL13^+^ Tregs maintain fundamental Treg characteristics.

**Figure 4.**
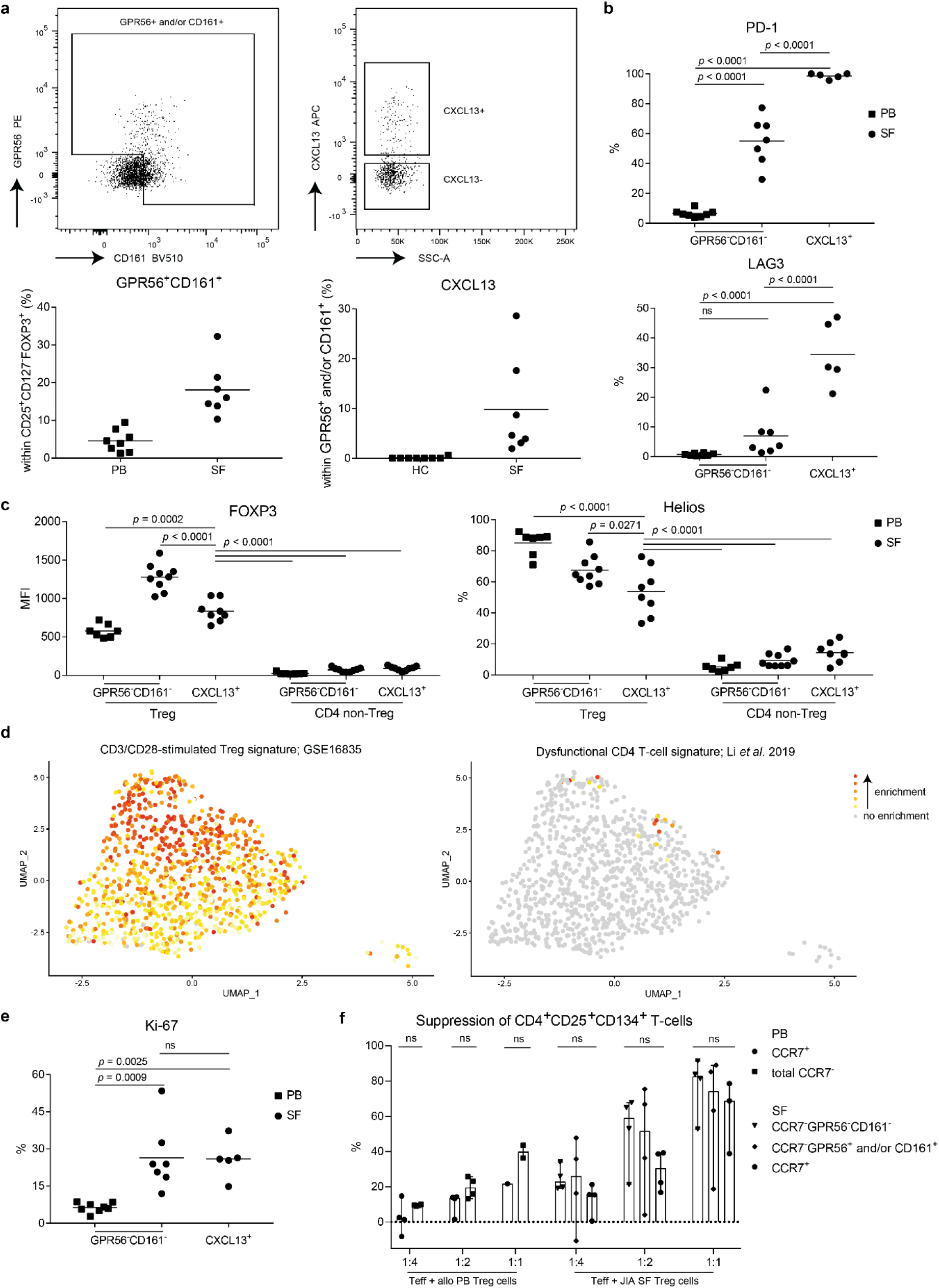
CD161^+^GPR56^+^CXCL13^+^ synovial fluid Tregs are highly differentiated and suppressive. **(a)** Representative gating (upper row) and quantification (lower row) of GPR56^+^ and/or CD161^+^ expression within CD127^low^CD25^high^FOXP3^+^ Tregs (left) and CXCL13 expression within this subset (left). Quantification is shown in control peripheral blood (PB) Tregs (*n* = 8) and synovial fluid (SF)-derived Treg from Juvenile Idiopathic Arthritis (JIA) patients (*n* = 7). **(b)** Quantification of PD-1 (upper) and LAG3 (lower) within control PB GPR56^-^CD161^-^ (*n* = 8), synovial fluid (SF) GPR56^-^CD161^-^ (*n* = 7) and SF CXCL13^+^ (GPR56^+^ and/or CD161^+^, and CXCL13^+^; *n* = 5) Tregs. **c** Quantification of Helios (left) and FOXP3 (right) within control PB GPR56^-^CD161^-^ (*n* = 7), SF GPR56^-^CD161^-^ (*n* = 9) and SF CXCL13^+^ (GPR56^+^ and/or CD161^+^, and CXCL13^+^; *n* = 8) Tregs and CD4 non-Tregs (CD127^+^CD25^low^FOXP3^-^). **(d)** Gene set analysis of a CD3/CD28 stimulated Treg (GSE16835) (left) and a dysfunctional CD4 T-cell signature^41^ (right). Enrichment is calculated per cell; grey signifies no enrichment and yellow to red shows increasing enrichment. **(e)** As per **(b)** for Ki-67. **(f)** Suppression of CD4 effector T-cells, assay as per Long et al.^31^. Sorted CD3^+^CD25^-^ T-cells (10.000) were cultured with Tregs (CCR7^+^ (cluster 1), CCR7^-^, CCR7^-^GPR56^-^CD161^-^ (cluster 2-3), CCR7^-^GPR56^+^ and/or CD161^+^ (cluster 4)) derived from control PB or SF of JIA patients in varying ratio’s (1:4, 1:2, 1:1) to quantify the suppression induced by Tregs. Control PB *n* = 4, SF *n* = 3-4. Statistical comparisons were performed using Friedman’s test. **(c, e, f)** Data are representative of two independent experiments. **(c** and **e)** Statistical comparisons were performed using one-way ANOVA with Tukey’s correction for multiple testing.

Genes associated with TCR stimulation showed the highest enrichment in clusters 3 and 4 (Figure 4d). Upregulation of TCR signaling was also reflected in the gene ontology analysis (Supplementary table 2) and their previously shown clonality (Figure 2a-c). Ongoing TCR stimulation has been associated with exhaustion, but exhaustion-associated genes were enriched in only 1.84% (18/980 Tregs) of the Tregs, and not particularly in cluster 4 (Figure 4d). Supporting the absence of exhaustion-associated gene signatures in cluster 4 is the high expression of Ki-67, a marker for recent proliferation, similar to other SF-derived Tregs (Figure 4e).

CD4^+^ non-Treg cells can transiently express FOXP3 upon activation, while maintaining effector functions^30^. To verify that our Tregs are *bona fide* suppressor cells, we assessed Treg-specific characteristics. On transcriptome level we observed higher expression of *BHLHE40, IFNG* and *ID2* in cluster 4 compared to the other SF Tregs, which are TFs associated with cytokine expression. However, on protein level we could not detect significant levels of IFNγ (1.63%), IL-2 (3.88%) or IL-17 (2.2%) produced by GPR56/CD161^+^ CXCL13^+^ SF Tregs (Supplementary figure 6a) indicating that these cells are true Tregs. In addition, we performed a suppression assay as developed by Long *et al*.^31^ to assess the suppressive capacity of cluster 4 Tregs. We sorted cluster 4 Tregs based on CCR7^-^GPR56 and/or CD161 expression (for gating strategy see Supplementary figure 6b), which includes CXCL13 expressing Tregs. In contrast, within the CD161^-^GPR56^-^ Treg population CXCL13^+^ Tregs comprised only 0.8% of the cells. In addition, we included CCR7^+^ Tregs for resting/quiescent Tregs and CCR7^-^ Tregs negative for both GPR56 and CD161 from both SF and control PB. Similar levels of CD4 effector T-cell suppression were observed by all SF Treg subsets (Figure 4f). These data show that Tregs from cluster 4 are proficient suppressor cells.

## Discussion

We show that even though Tregs in SF of JIA patients obtain a dominant Th1 co-transcriptional program, substantial heterogeneity can be observed. The largest proportion of SF Tregs actually comprise of recently arrived cells (*CCR7, KLF2, TCF7, LEF1*) which seem to differentiate to three activated (e)Treg phenotypes that are not completely discrete but form a gradient of transcriptional states. Two of these phenotypes are more classical eTregs characterized by *CTLA4, TIGIT, GZMA, TNFRSF8, CCR5*, and *TBX21*, whereas the third activated Treg cluster contains Tregs expressing *CXCL13, GPR56*, and *KLRB1*. Tregs from the latter cluster bear a more distinct TCR profile, while the eTregs of clusters 2 and 3 are more closely related. Lastly, next to BATF as shared regulator in eTreg differentiation, the here identified Treg regulators BHLHE40 and JAZF1 regulate primarily genes associated with cluster 4 and clusters 2/3 Tregs, respectively. The TF BHLHE40 (also known as DEC1 or BHLHB2) is upregulated upon TCR activation. BHLHE40 in Th-cells is associated with a pro-inflammatory phenotype^32,33^; however, in Tregs this TF seems crucial for long-term maintenance of the Treg pool, and adoptive transfer of BHLHE40 expressing Tregs in a colitis model in mice prevents wasting disease^34^. JAZF1 (TIP27) has been mostly studied for its role in gluconeogenesis and lipid metabolism in relation to the development of type 2 diabetes mellitus, and its signaling also results in decreased expression of proinflammatory cytokines^35,36^. Our data suggest that JAZF1, and its downstream target genes, skew Treg metabolism in the chronic inflammatory setting, but this remains to be fully elucidated. These data support the notion that Tregs within SF follow different and unique routes of adaptation.

In both mice and humans, studies have explored PB to tissue Treg differentiation. However, data on functional adaptation pathways of Tregs in inflamed tissues is lacking. Here we show that infiltrating Tregs are heterogeneous in chemokine receptor expression upon migration to SF indicating (partial) unspecific homing in response to inflammatory signals, with local cues and antigen(s) inducing preferential differentiation to, and expansion of, Th1-skewed Tregs. The acquisition of a Th-associated co-transcriptional program is crucial for Treg survival and function in those inflammatory environments^37^. The demonstrated heterogeneity also suggests that upon local cues Tregs can induce subtle shifts in phenotype, possibly to suppress immune responses under changing inflammatory conditions. Especially for cluster 2 and 3 Tregs this seems likely since they share many clonotypes but show phenotypic differences in functional Treg markers. For GPR56^+^CD161^+^CXCL13^+^ Tregs (cluster 4), however, clonotypes are less overlapping suggesting a more predetermined route of these Tregs, perhaps based on TCR-specificity or affinity, when migrating to the site of inflammation. Thereby the question rises where the imprinting of Treg differentiation occurs. Chemokine receptor expression is determined in secondary lymphoid structures and functional marker heterogeneity seems to be partially predetermined in the periphery^38–40^. Our data suggests that there is further extensive local plasticity and differentiation, which is supported by the observation that different T-cell subsets show a similar phenotypical adaptation in local inflammatory environments.

Several scRNA-sequencing studies in autoimmune and tumor settings have shown presence of CD4 and CD8 T-cell subsets characterized by CXCL13 and inhibitory immune checkpoints^16,41–44^ indicating a partial mirrored phenotype amongst inflammation-derived T-cells. However, for FOXP3^+^ Tregs this phenotype hasn’t been described before. These CXCL13^+^ subsets express high levels of PD-1 and other inhibitory immune checkpoints, proliferate, are clonally expanded and have lost classical CD4 and CD8 T-cell effector functions including cytokine production^16,41–44^. These cells have a T peripheral helper-like phenotype with respect to CXCL13 and PD-1 expression as well as absence of CXCR5 and Bcl6^43^, but do not produce IL-21. Genes such as *PDCD1, CCL5, GNLY, HAVCR2* and *CCL4* expressed by these cells suggest a terminal stage of differentiation. However, Li *et al*.^41^ showed that these dysfunctional cells are tumor-reactive, and gaining CXCL13 expression might indicate acquisition of novel functions^44^. TCR stimulation, presence of inflammatory cytokines including type I interferons and especially TGFβ are known to induce CXCL13 expression^45^, which are all abundant in inflammatory environments. That these cells are not found in the periphery indicates that CXCL13 expression is acquired locally. CXCL13 expression suggests a role in ectopic lymphoid structure (ELS) formation as CXCL13 is a known B-cell attractant. Although, Tregs can regulate B-cell maturation within lymphoid structures it is unknown if they can also help forming ELS via CXCL13 expression. Supportive hereof could be expression of extracellular matrix organization-related genes, such as *ADAM19, COL6A3, COL9A2, CTSH, CCL4*, and *CCL5* by a fraction of these CXCL13^+^ Tregs. Expression of these genes could also indicate an involvement in tissue repair via extracellular matrix organization after inflammation-related damage in JIA patients^46^. Furthermore, in a murine model tissue repair recently has been shown as a function of Tregs at the site of myocardial infarctions^47^. Additional markers expressed by CXCL13^+^ cells, such as GPR56 for SF Tregs, seem more cell-specific since they are not co-expressed in CXCL13^+^ (SF) non-Treg. Furthermore, for SF GPR56^+^CD161^+^ Tregs (containing CXCL13^+^ Tregs) there is no loss of classical functioning as these Tregs remain proficient suppressors. This indicates that CXCL13 expression and expression of inhibitory immune checkpoints is likely dependent on the T-cell subset concerned. Whether these CXCL13^+^ T-cell subsets, and specifically CXCL13^+^ Tregs, in the autoimmune setting are beneficial or pathogenic needs to be elucidated.

Limitations of this study include the risk of contamination of the sorted Tregs with non-Tregs. However, CD127^high^CD25^low^ Tregs were sorted with a purity >98% with a FOXP3 intracellular staining confirming >90% FOXP3 expression and 96.7% of the Tregs contained at least one FOXP3 transcript. Another concern could be the sample size of 3 JIA patients for the scRNA-sequencing. Therefore, we confirmed our findings in a public 10x Genomics sequenced SF T-cell dataset including Tregs. We could also confirm presence of CXCL13^+^ Tregs at protein level further supporting our transcriptomic findings. Furthermore, there is still a limited understanding of gene signatures representing functions or states in human Tregs. For example, a definition of exhausted Tregs is lacking with many of the genes defined for CD8 and CD4 T-cells actually signifying enhanced Treg activity such as *PDCD1, ENTPD1* (CD39), and *LAG3*. It is of importance to further explore (human) Tregs in inflammatory settings to improve our understanding of their (dys)function.

In recent years, several Treg-targeted therapeutic strategies have been implemented in the clinic or are in clinical trial including inhibitory immune checkpoints such as CTLA-4 and LAG3^4^. Our data suggest that not all Tregs will be targeted equally since expression of inhibitory immune checkpoints differs amongst Tregs in one environment. It would be valuable to study patient inter- and intra-variability (if heterogeneity is dynamic) regarding Treg heterogeneity since this could aid in improving personalized Treg-based therapeutic strategies; is for example PD-1, LAG3 or a combination better suited for a patient?^48^ Additionally, when aiming to prevent adverse side-effects it might be more specific to target chemokine receptors such as CCR4 in clinical trial^4^ because expression of this chemokine receptor depends on the environment.

In conclusion, our study reveals a heterogeneous population of Tregs at the site of inflammation in JIA. SF Treg differentiate to a classical eTreg profile with a more dominant suppressive or cytotoxic profile that share a similar TCR repertoire, or towards GPR56^+^CD161^+^CXCL13^+^ Tregs with a more distinct TCR repertoire. The latter cluster of Tregs is also mirrored in other T-cell subsets at the site of inflammation. Finally, the novel Treg regulon BHLHE40 seems to drive differentiation towards primarily GPR56^+^CD161^+^CXCL13^+^ Tregs and JAZF1 towards the classical eTreg phenotype.

## Methods

### Patient samples

For scRNA-sequencing, patients with oligo JIA (*n* = 3, 3/3 female) were enrolled in the pediatric rheumatology department at the University Medical Center of Utrecht (the Netherlands). The average age was 9.3 years (range 7-12 years) with a disease duration at the time of inclusion of 7 years (range 4-10 years). Two patients were without medication, and one received methotrexate maintenance therapy. HLA-B27 status has not been assessed. For flow cytometry, patients with oligo JIA (*n* = 17; 40% female, average age 14.2 [2-19 years]) of which 30% had extended and 55% persistent oligo-articular disease were included. Ten patients were without medication, 5 on methotrexate, 2 on NSAIDs and 3 on anti-TNF maintenance therapy. In addition, PB of controls (*n* = 21; 58% female, average age 39.3 [25-62 years]) were included.

Active disease was defined by physician global assessment of ≥ 1 active joint (swelling, limitation of movement), and inactive disease was defined as the absence hereof. During an outpatient clinic visit, SF was obtained by therapeutic joint aspiration of the affected joints, and blood was withdrawn via vein puncture or an intravenous drip catheter. The study was conducted in accordance with the Institutional Review Board of the University Medical Center Utrecht (approval no. 11-499/C). PB from healthy adult volunteers was obtained from the Mini Donor Service at University Medical Center Utrecht. The research was carried out in compliance with the Declaration of Helsinki. Informed consent was obtained from all the participants and/or from their parents/guardians/legally authorized representatives.

SF of JIA patients was incubated with hyaluronidase (Sigma-Aldrich) for 30 min at 37°C to break down hyaluronic acid. Synovial fluid mononuclear cells (SFMCs) and peripheral blood mononuclear cells (PBMCs) were isolated using Ficoll Isopaque density gradient centrifugation (GE Healthcare Bio-Sciences, AB).

### Single-cell mRNA-sequencing

Live CD3^+^CD4^+^CD25^+^CD127^low^ cells were sorted from fresh SF (Supplementary figure 1a) into 384-well hard shell plates (Biorad) with 5 μl of vapor-lock (QIAGEN) containing 100-200 nl of RT primers, dNTPs and synthetic mRNA Spike-Ins and immediately spun down and frozen to -80°C. Cells were prepared for SORT-seq as previously described^49^. Illumina sequencing libraries were then prepared with the TruSeq small RNA primers (Illumina) and sequenced single-end at 75 basepair read length with 60.000 reads per cell on a NextSeq500 platform (Illumina). Sequencing reads were mapped against the reference human genome (GRCh38) with BWA.

### Single-cell mRNA-sequencing analysis

Quality control was performed in R with the scater package v1.12.2^50^ and cells were dropped when the number of genes, number of UMI’s and/or the percentage of mitochondrial genes was over 3 median absolute deviations under/above the median. Afterwards principal component analysis (PCA)-outliers were removed with the package mvoutlier v1. The raw data expression matrices were subsequently analyzed using Seurat v2-4^51–53^ following the outline provided by the distributor (https://satijalab.org/seurat/). Each dataset was normalized and the cell-cycle was regressed out using SCTransform^54^. Thereupon, the SF datasets were integrated with PrepSCTIntegration followed by FindIntegrationAnchors and IntegrateData for SCTransform-processed data.

For dimensionality reduction first the principal components (PCs) were calculated (RunPCA) and clustering was performed with UMAP (RunUMAP: 30 dimensions; FindNeighbors: clustering resolution of 1). One cluster was removed from further analyses (Supplementary figure 2a) since it was of ambiguous origin (e.g. hybrid, transferred extracellular vesicles or doublets). Subsequent differential gene expression was performed using the MAST test (standard settings) with a p-adjusted value < 0.05 considered statistically significant. For visualization the functions DoHeatmap, Dimplot, Featureplot and plot_density were employed. UMAPs were plotted with raw mRNA counts or using the new nebulosa algorithm^55^ based on kernel density estimation to handle sparsity of scRNA-sequencing data.

Gene set enrichment analysis was performed with gene sets derived from the literature (Ferraro *et al*.^13^ for a human Treg signature, Niedzielska *et al*.^56^ for a shared tissue Treg signature, De Simone *et al*.^57^ for a tumor-infiltrating Treg signature, and Li *et al*.^41^ for a dysfunctional signature of CD4^+^ T-cells) or the MSigDB C7 database (GSE61077 for CD44^hi^CD62L^lo^ versus CD44^lo^CD62L^hi^ murine Tregs, GSE11057 for naive T-cells versus effector memory human CD4 T-cells, GSE16835 for CD3/CD28 stimulated versus *ex vivo* Treg) with AUCell^14^. The cut-off for enrichment of a gene set in a cell was defined using the AUC. Per cluster the proportion of cells enriched for the gene set was calculated and compared with the Chi-square test. Gene Ontology pathway analyses implementing the probability density function were performed using ToppFun (https://toppgene.cchmc.org/enrichment.jsp) with as input the differentially expressed genes belonging to each Treg cluster, with a false discovery rate (FDR)-corrected p-value < 0.05 defining significance.

For pseudotime trajectory analysis Monocle v3^17^ was used with a trajectory predicted using standard settings based on the clustering previously performed with Seurat. The principal root node was estimated mathematically with the function get_earliest_principal_node. Network inference analysis employing SCENIC^14^ was performed in python v3.6 with Jupyter Notebook v6.1.5 using the UMAP clustering as starting point to define regulons. In short, co-expression based on the raw count data and DNA motif analysis is used to obtain transcription factors and their target genes using standard settings. Activity of these potential TFs and targets (regulons) are analyzed per cell and finally the scores per cell were combined to compare clusters with each other to define regulators that might drive differentiation within the SF environment. The scRNA-sequencing count data generated for this study have been submitted to a public repository on GitHub (https://github.com/lutterl/JIA-synovial-fluid-Tregs-scRNAseq). Raw data files will be made publicly available before publication.

### Single cell TCR-sequencing analysis

RNA- and TCR-sequencing profiling data of single cell SF Tregs employing 10X genomics were downloaded from GSE160097 [<https://www.ncbi.nlm.nih.gov/geo/query/acc.cgi?acc=GSE160097]>. TCR-sequencing data was analyzed using scRepertoire following the outlined guidelines^58^. In short, T-cell receptor alpha locus (TRA) and TCR beta locus (TRB) data was combined based on the cell barcode. If there was more than one TRA and/or TRB chain detected the most prevalent chain was selected for integration. RNA-sequencing and TCR-sequencing data was combined and subsequent T-cell clustering was performed with Seurat as described above. TCR-sequencing analysis was performed on the nucleotide level.

### Flow cytometry

#### Immunophenotyping

PBMCs and SFMCs were thawed and resuspended in RPMI1640 (Gibco) supplemented with 10% Fetal Bovine Serum (FBS). For measurements of CXCL13 the cells were cultured for 5 hours at 37°C in RPMI supplemented with 10% human AB serum and GolgiStop (1/1500; BD Biosciences). For cytokine measurements the cells were plated in the presence of anti-CD3/CD28 (Dynabeads^®^ Human T-activator CD3/CD28, ThermoFisher Scientific) at a 1:5 ratio (bead:cell) at 37°C. After 19 hours cells were incubated for 5 hours with GolgiStop. After stimulation, cells were stained with surface antibodies for 20 min at 4°C. The following antibodies were used: fixable viability dye eF780 or eF506 (eBioscience), anti-human CD3 AF700 (clone UCHT1), GPR56 PE-Cy7 (clone CG4), PD-1 PerCP-Cy5.5 (clone EH12.2H7; Biolegend), CD4 BV785 (clone OKT4; eBioscience), CD25 BV711 (clone 2A3), CD161 BV510 (clone DX12), CD161 PE-Cy5 (clone DX12), Helios PE (clone 22F6; BD), CD127 BV605 (clone A019D5; Sony Biotechnology), and LAG-3 PE (polyclonal; R&D). For intranuclear/cellular staining the Intracellular Fixation & Permeabilization Buffer Set (eBioscience) was used, and staining was performed for 30 minutes at 4°C. The following antibodies were used: Ki-67 FITC (clone mip1; Dako), CXCL13 APC (clone 53610; R&D), IFNγ PerCP-Cy5.5 (clone 4S.B3), FOXP3 eF450 (clone PCH101), IL-17 FITC (clone eBio64DEC17; eBioscience), FOXP3 PE-CF594 (clone 259D/C7), IL-2 PE (clone MQ1-17H12; BD). Data acquisition was performed on a BD LSRFortessa (BD Biosciences) and data were analyzed using FlowJo Software v10 (Tree Star Inc.).

#### Suppression assay

Suppression assays were performed according to the outline provided by Long *et al*.^31^. In short, effector cells (live CD3^+^CD25^-^, lowest 50% of CD25 stained CD3^+^ T-cells) and Tregs (CD3^+^CD4^+^CD127^low^CD25^high^ subdivided into CCR7^+^, CCR7^-^GPR56^-^CD161^-^ (DN) and CCR7^-^GPR56^+^ and/or CD161^+^) were isolated from frozen PBMC and SFMC using the FACS Aria III (BD). Both PBMC and SFMC were ‘rested’ for 24 hours in RPMI1640 supplemented with 10% FBS at 37°C prior to sorting. Antibodies used for sorting were: CD3 AF700 (clone UCHT1), CD127 AF647 (clone HCD127), GPR56 PE-Cy7 (clone CG4; Biolegend), CD25 BV711 (clone 2A3), CD161 BV510 (clone DX12; BD), CD4 FITC (clone RPA-T4), and CCR7 PE (clone 3D12; eBioscience). Effector cells were labeled with 2µM ctViolet (Thermo Fisher) and cultured alone or with different ratios of sorted Tregs (1:1, 1:2, 1:4). Cells were cultured in RPMI1640 media containing 10% human AB serum with addition of L-Glutamine and Penicillin/Streptomycin. Effector cells were stimulated by anti-CD3/CD28 dynabeads at a 1:28 ratio (ThermoFisher Scientific), and incubated for 48 hours at 37°C. Read out on a BD LSRFortessa (BD Biosciences) was performed using Fixable viability dye eF506 (eBioscience), ctViolet (Thermo Fisher), CD8 PE-Cy7 (clone SK1; BD), CD4 BV785 (clone OKT4), CD25 BV711 (clone 2A3; eBioscience), CD3 AF700 (clone UCHT1) and CD134 PerCP-Cy5.5. (OX40, clone Ber-ACT35; Biolegend), with CD25 and CD134 as surrogate markers for proliferation^31^. Data were analyzed using FlowJo Software v10 (Tree Star Inc.).

### Statistical analysis

Statistical analyses were performed with Pearson’s Chi-squared test, a Friedman’s test with Dunn’s post hoc, or a one-way ANOVA with Tukey’s post-hoc test if applicable. Paired data comparisons with missing values were analyzed with a mixed-effect model (Restricted Maximum Likelihood). Analyses were performed in Graphpad Prism v7.04 and v8.3, Excel Office v2017 and R v3.5.2-4.1.0.

## Acknowledgements

We would like to thank Sytze de Roock for help with selecting donor samples, Single Cell Discoveries for running our single cell RNA-sequencing (SORT-seq), and Pawel Durek for providing us with the single cell TCR-sequencing data. E.C. Brand was supported by the Alexandre Suerman program for MD and PhD candidates of the University Medical Centre Utrecht, Netherlands. F. van Wijk is supported by a VIDI grant from the Netherlands Organization for Scientific Research (ZonMw, 91714332). E.C. Brand and F. van Wijk are co-applicants on an Investigator Initiated research grant of Pfizer unrelated to this manuscript.

## Author contributions

Conceptualization: L.L., F.v.W. Patient selection and clinical interpretation: L.L., B.V. Performed experiments: L.L., M.vd.W. Data analysis: L.L., M.vd.W., E.C.B. Supplied scTCR-sequencing dataset: P.M., M.M. Writing: L.L., M.v.d.W, J.v.L., F.v.W. Supervision: F.v.W. All authors reviewed and edited the manuscript.

## Notes

**Conflict of interest** The authors have declared that no conflict of interest exists.

### Competing Interest Statement

The authors have declared no competing interest.

