## Supplementary Figures for "Human regulatory T-cells locally differentiate and are functionally heterogeneous within the inflamed arthritic joint"

### Supporting information Lutter et al

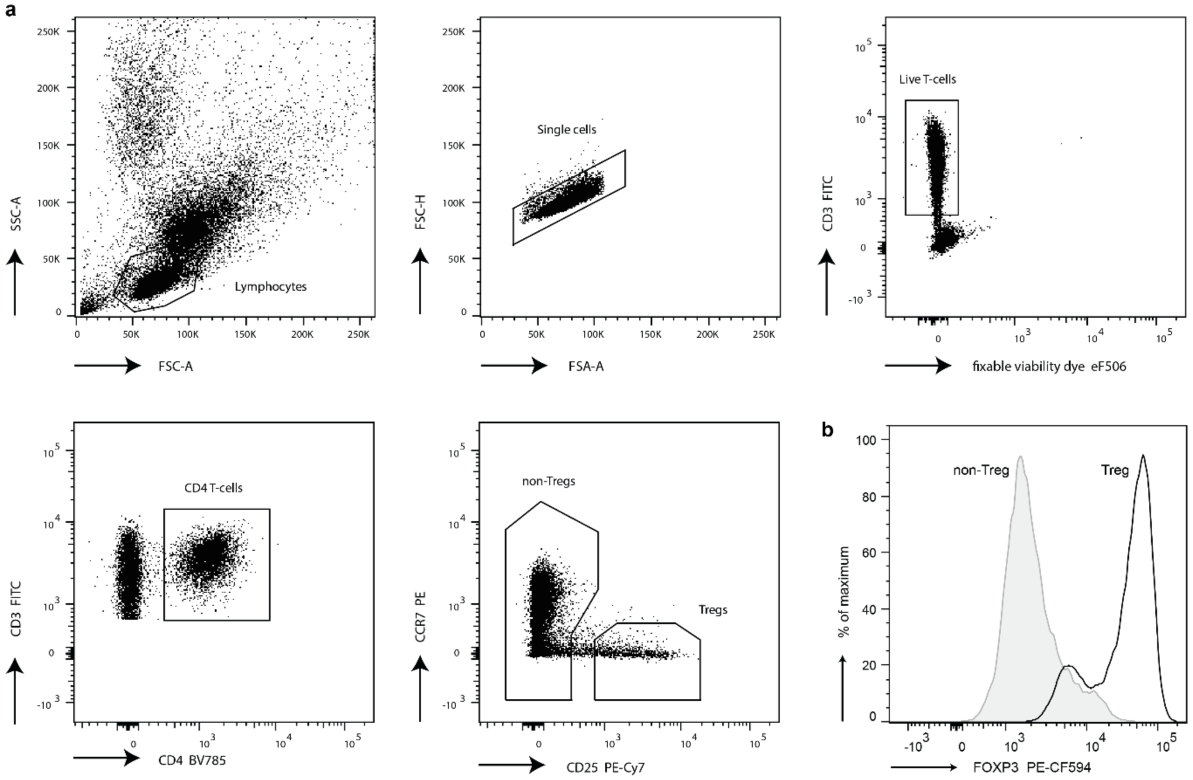


**Supplementary figure 1.** Gating strategy of synovial fluid derived Tregs. **(a)** Representative gating strategy to sort Tregs derived from synovial fluid (SF). **(b)** Representative FOXP3 median fluorescence.

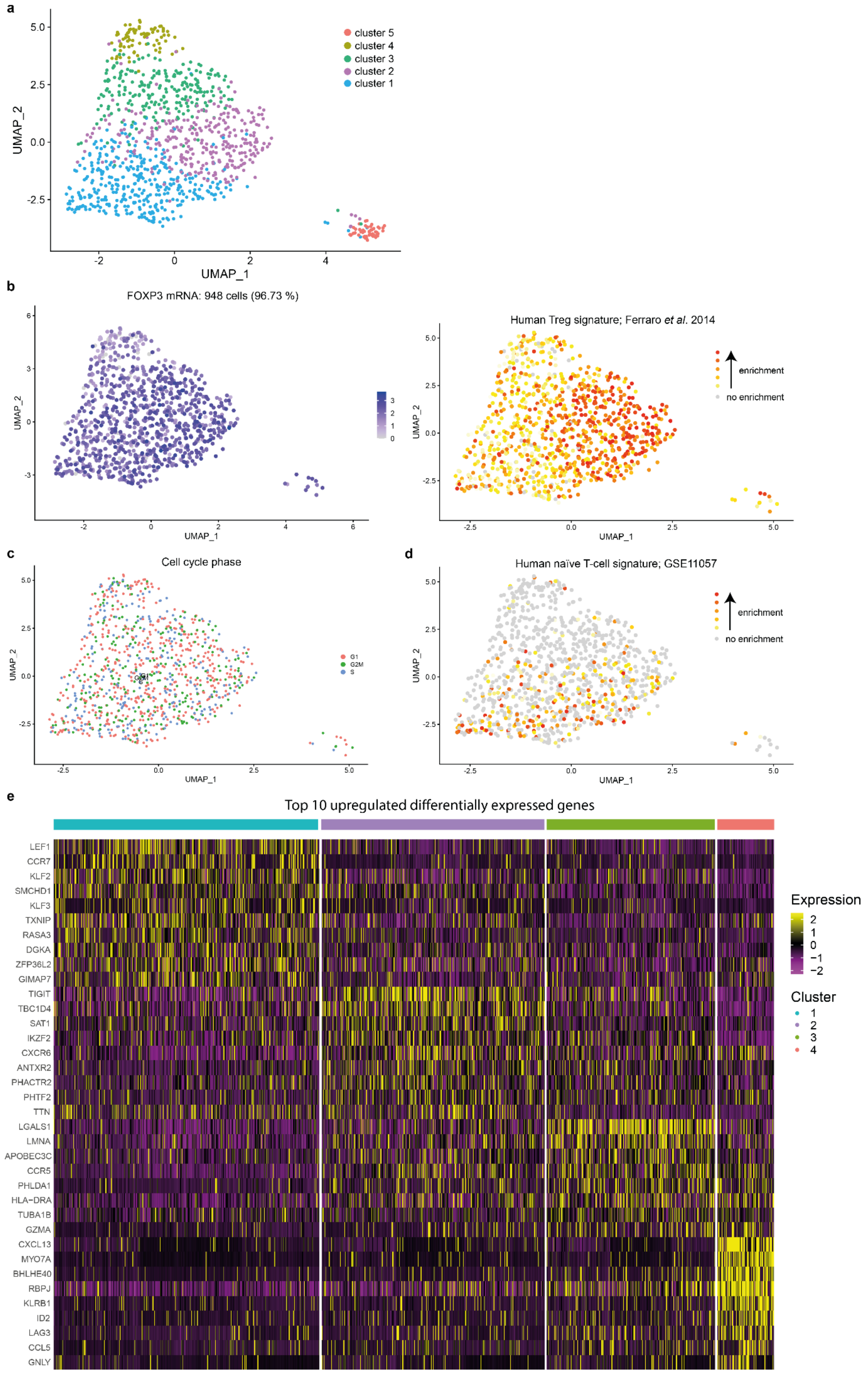
a
UMAP_1
UMAP_2
cluster 5
cluster 4
cluster 3
cluster 2
cluster 1
b
FOXP3 mRNA: 948 cells (96.73 %)
Human Treg signature; Ferraro et al. 2014
enrichment
no enrichment
c
Cell cycle phase
G1
G2M
S
d
Human naïve T-cell signature; GSE11057
enrichment
no enrichment
e
Top 10 upregulated differentially expressed genes
LEF1
CCR7
KLF2
SMCHD1
KLF3
TXNIP
RASA3
DGKA
ZFP36L2
GIMAP7
TIGIT
TBC1D4
SAT1
IKZF2
CXCR6
ANTXR2
PHACTR2
PHTF2
TTN
LGALS1
LMNA
APOBEC3C
CCR5
PHLDA1
HLA−DRA
TUBA1B
GZMA
CXCL13
MYO7A
BHLHE40
RBPJ
KLRB1
ID2
LAG3
CCL5
GNLY
Expression
Cluster

**Supplementary figure 2.** Heterogeneity and phenotypical profile of synovial fluid Tregs. **(a)** Dimensionality reduction (UMAP) of all sequenced Tregs that passed the quality control. Cluster 5 is of ambiguous composition (e.g. hybrid, transferred extracellular vesicles or doublets) and not included in further analysis. **(b)** mRNA FOXP3 expression (left) and gene set enrichment of a human core Treg signature[19] (right). Grey means no FOXP3 mRNA expression or Treg signature enrichment, with in blue cells with FOXP3 mRNA (left), and yellow to red increasing shows enrichment of the Treg signature (right). **(c)** UMAP with the cells colored by phase of the cell cycle; pink = G1, green = G2M, blue = S. Pearson's Chi-squared test revealed no association between the cell cycle phase and the clustering ($p$ = 0.8512). **(d)** Heatmap of the top 10 upregulated differentially expressed genes, based on $\log_2$ fold change, per cluster. Expression ranges from purple (no raw count mRNA expression) to yellow (presence of mRNA expression of the concerned gene). **(e)** Similar to **(b)** but for a gene set upregulated in naïve compared to effector memory human CD4 T-cells (GSE11057).

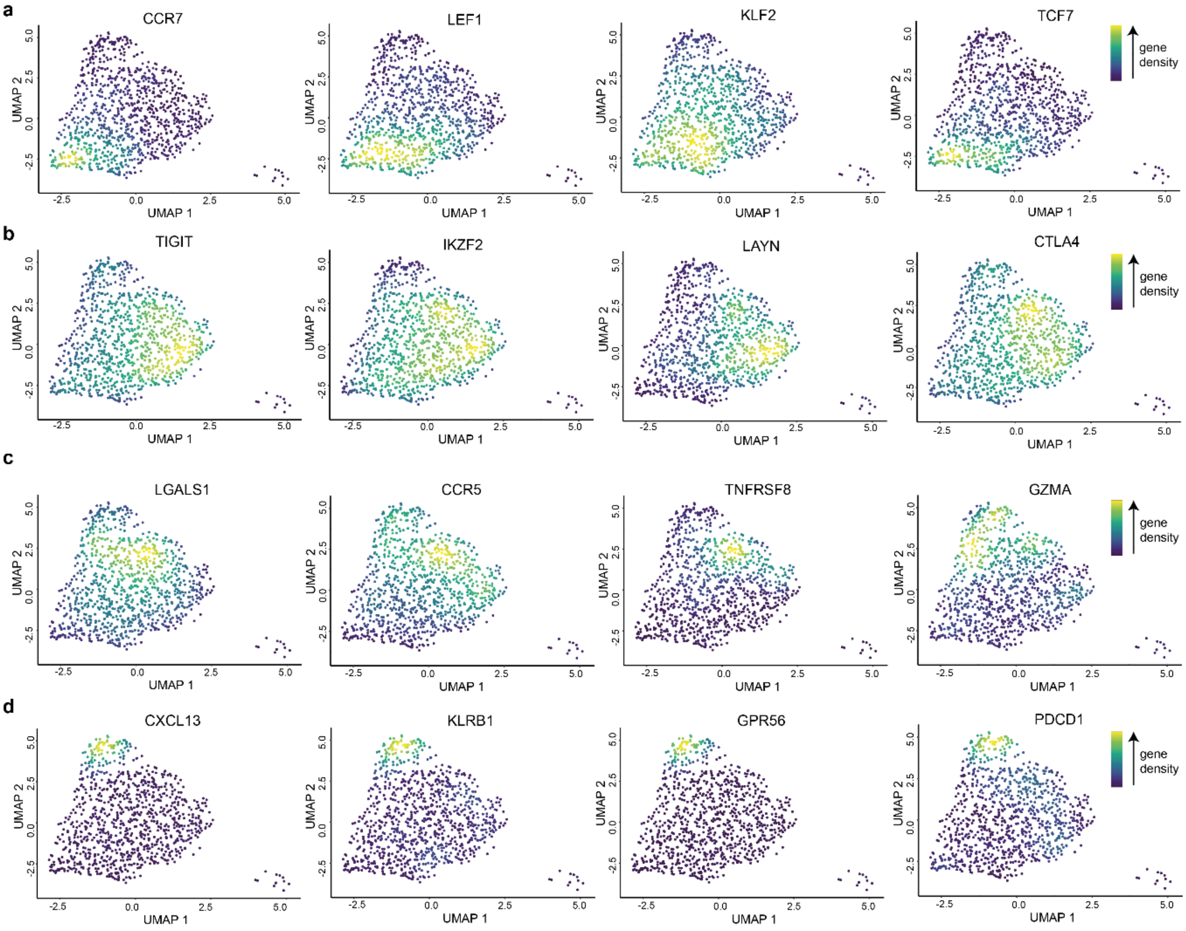


**Supplementary figure 3.** Synovial fluid Treg cluster-identifying genes. **(a)** UMAPs showing the nebulosa density of the selected differentially expressed genes *CCR7*, *LEF1*, *KLF2*, and *TCF7* for cluster 1 Tregs. The nebulosa density (based on kernel density) ranges from blue to yellow, with in yellow the highest kernel density, thus the highest (estimated) expression of the shown gene. **(b)** Similar to **(a)** but for *TIGIT*, *IKZF2* (encoding Helios), *LAYN*, and *CTLA4* which are differentially expressed by cluster 2 Tregs. **(c)** Similar to **(a)** but for *LGALS1*, *CCR5*, *TNFRSF8*, and *GZMA* which are differentially expressed by cluster 3 Tregs. **(d)** Similar to **(a)** but for *CXCL13*, *KLRB1*, *GPR56*, and *PDCD1* which are differentially expressed by cluster 4 Tregs.

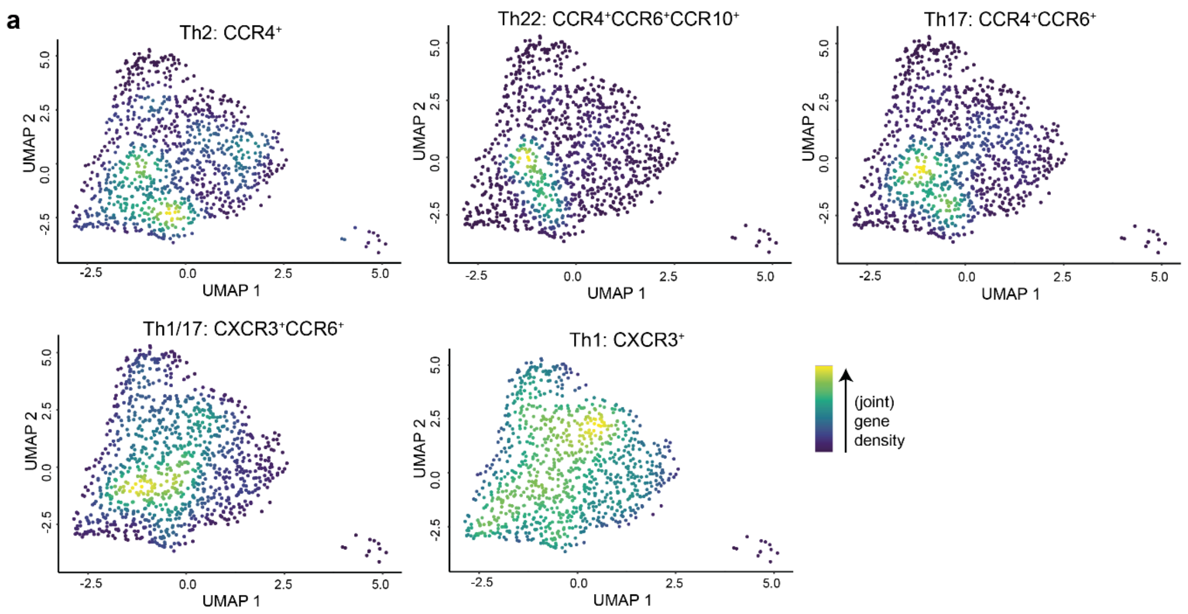


**Supplementary figure 4.** Preferential expansion of Th1-skewed Tregs in synovial fluid. **(a)** UMAPs showing the nebulosa density of chemokine receptor gene expression combinations (positive expression) that are associated with Tregs that have obtained a T helper (Th)-type co-transcriptional program. Upper row from right to left: *CCR4* for Th2-skewed Tregs, *CCR6*, *CCR4* and *CCR10* expression for Th22-skewed Tregs, and *CCR6* and *CCR4* expression for Th17-skewed Tregs. Lower row from right to left: *CXCR3* and *CCR6* for Th1/Th17-skewed Tregs and *CXCR3* for Th1-skewed Tregs. The nebulosa density (based on kernel density) ranges from blue to yellow, with in yellow the highest kernel density, thus the highest (estimated) expression of the combination of genes.

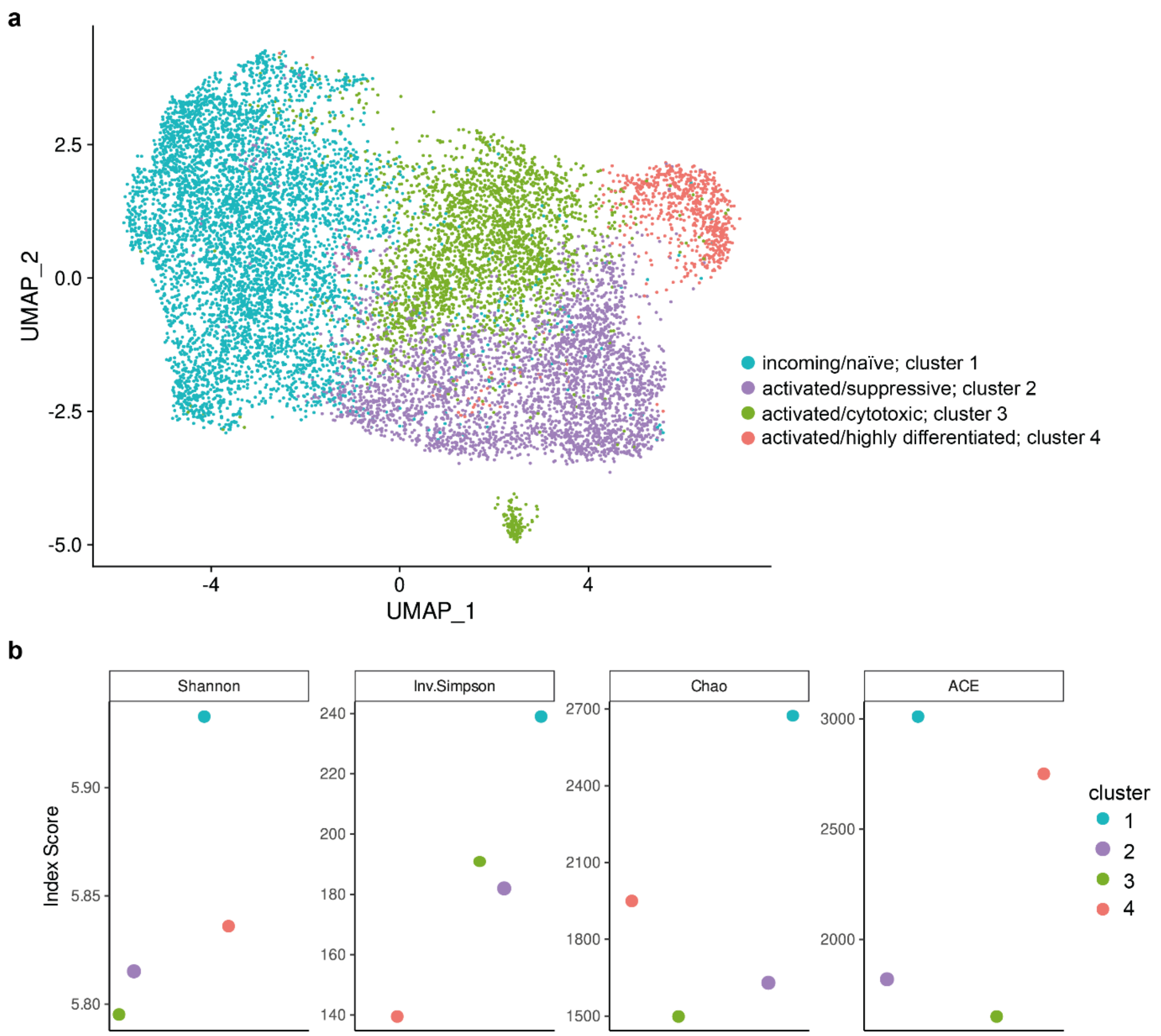


**Supplementary figure 5.** Clonal expansion and clonal overlap of synovial fluid-derived Tregs. **(a)** Dimensionality reduction (UMAP) similar to Fig. 1a but of a 10X genomics sequenced synovial fluid (SF)-derived Treg dataset (GSE160097) with the cells colored by cluster. Colors are identical to the clustering of Fig. 1a. **(b)** Morisita, inverse Simpson, Chao and ACE diversity indexes calculated per SF Treg cluster. Each index is a measure for the diversity of the T-cell receptor repertoire per cluster with higher values indicating less diversity of the repertoire.

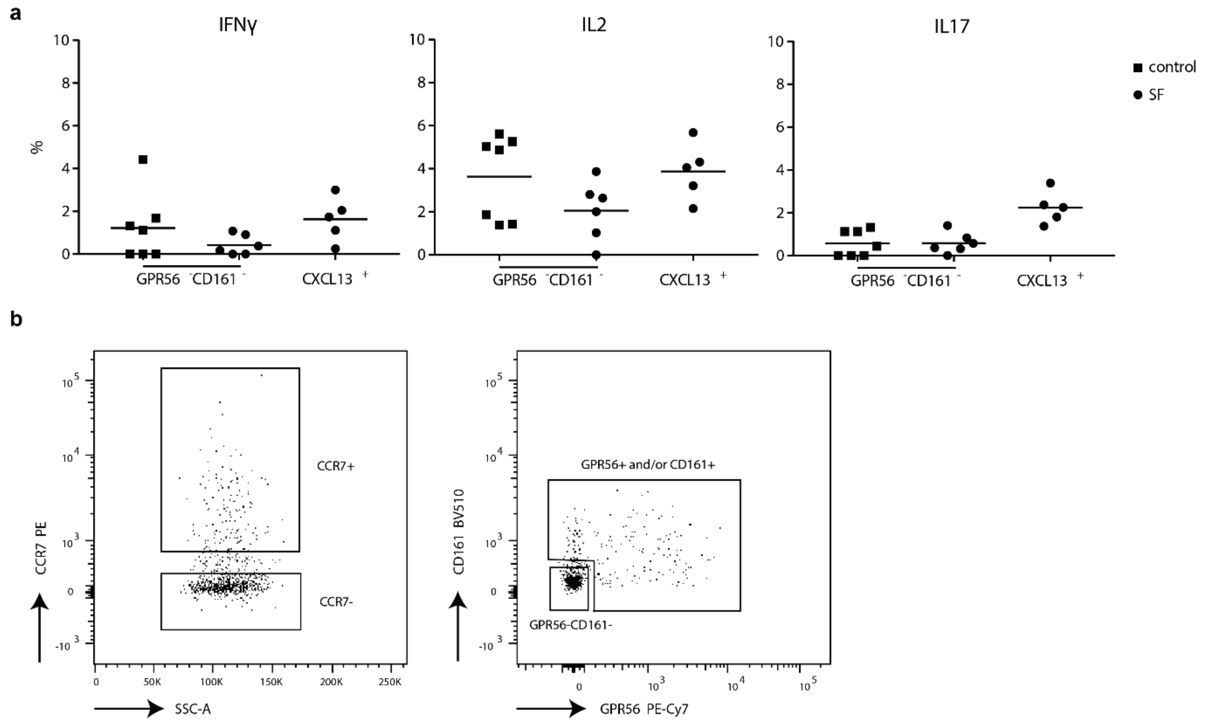


**Supplementary figure 6** $CD161^+GPR56^+CXCL13^+$ synovial fluid Tregs are highly differentiated and suppressive cells. **(a)** Percentages of IFNγ (left), IL-2 (middle) and IL-17 (right) expression within control peripheral blood (PB) $GPR56^-CD161^-$, synovial fluid (SF) $GPR56^-CD161^-$ and SF $CXCL13^+$ ($GPR56^+$ and/or $CD161^+$, and $CXCL13^+$) Tregs ($CD127^{low}CD25^{high}FOXP3^+$). control PB $n = 7$, SF $n = 6$. Data are representative of two independent experiments. **(b)** Representative gating strategy to sort $CCR7^+$, $CCR7^-$ (in PB), $CCR7^-GPR56^-CD161^-$ (in SF) and $CCR7^-GPR56^+$ and/or $CD161^+$ (in SF) Tregs for the suppression assay (gating of Tregs as per Supplementary figure 1a).
